# Contrasting patterns of evolutionary constraint and novelty revealed by comparative sperm proteomic analysis

**DOI:** 10.1101/144089

**Authors:** Emma Whittington, Desiree Forsythe, Timothy L. Karr, James R. Walters, Steve Dorus

## Abstract

**Background:** Rapid evolution is a hallmark of reproductive genetic systems and arises through the combined processes of sequence divergence, gene gain and loss, and changes in gene and protein expression. While studies aiming to disentangle the molecular ramifications of these processes are progressing, we still know little about the genetic basis of evolutionary transitions in reproductive systems. Here we conduct the first comparative analysis of sperm proteomes in Lepidoptera, a group that broadly exhibits dichotomous spermatogenesis, in which males simultaneously produce a functional fertilization-competent sperm (eupyrene) and an incompetent sperm morph lacking DNA (apyrene). Through the integrated application of evolutionary proteomics and genomics, we characterize the genomic patterns associated with the origination of this unique spermatogenic process and assess the importance of genetic novelty in Lepidoptera sperm biology.

**Results:** Comparison of the newly characterized Monarch butterfly (*Danaus plexippus*) sperm proteome to those of the Carolina sphinx moth (*Manduca sexta*) and the fruit fly (*Drosophila melanogaster*) demonstrated conservation at the level of protein abundance and post-translational modification within Lepidoptera. In contrast, comparative genomic analyses across insects reveals significant divergence at two levels that differentiate the genetic architecture of sperm in Lepidoptera from other insects. First, a significant reduction in orthology among Monarch sperm genes relative to the remainder of the genome in non-Lepidopteran insect species was observed. Second, a substantial number of sperm proteins were found to be specific to Lepidoptera, in that they lack detectable homology to the genomes of more distantly related insects. Lastly, the functional importance of Lepidoptera specific sperm proteins is broadly supported by their increased abundance relative to proteins conserved across insects.

**Conclusions:** Our results suggest that the origin of heteromorphic spermatogenesis early in Lepidoptera evolution was associated with a burst of genetic novelty. This pattern of genomic diversification is distinct from the remainder of the genome and thus suggests that this transition has had a marked impact on Lepidoptera genome evolution. The identification of abundant sperm proteins unique to Lepidoptera, including proteins distinct between specific lineages, will accelerate future functional studies aiming to understand the developmental origin of dichotomous spermatogenesis and the functional diversification of the fertilization incompetent apyrene sperm morph.

## Introduction

Spermatozoa exhibit an exceptional amount of diversity at both the ultrastructure and molecular levels despite their central role in reproduction [1]. One of the least understood peculiarities in sperm variation is the production of heteromorphic sperm via dichotomous spermatogenesis, the developmental process where males produce multiple distinct sperm morphs that differ in their morphology, DNA content and/or other characteristics [2]. Remarkably, one sperm morph is usually fertilization incompetent and often produced in large numbers; such morphs are commonly called “parasperm”, in contrast to fertilizing “eusperm” morphs. Despite the apparent inefficiencies of producing sperm morphs incapable of fertilization, dichotomous spermatogenesis has arisen independently across a broad range of taxa, including insects, brachiopod molluscs and fish. This paradoxical phenomenon, where a substantial investment is made into gametes that will not pass on genetic material to the following generation, has garnered substantial interest, and a variety of hypotheses regarding parasperm function have been postulated [3]. In broad terms, these can be divided into three main functional themes: (**1**) facilitation, where parasperm aid the capacitation or motility of eusperm in the female reproductive tract, (**2**) provisioning, where parasperm provide nutrients or other necessary molecules to eusperm, the female or the zygote and (**3**) mediating postcopulatory processes, where parasperm may serve eusperm either defensively or offensively by delaying female remating, influencing rival sperm, or biasing cryptic female choice. Despite experimental efforts in a number of taxa, a robust determination of parasperm function has yet to be attained.

Dichotomous spermatogenesis was first identified in Lepidoptera [4], the insect order containing butterflies and moths, over a century ago and is intriguing because the parasperm morph (termed apyrene sperm), is anucleate and therefore lacks nuclear DNA. Although it has been suggested that apyrene sperm are the result of a degenerative evolutionary process, several compelling observations suggest that dichotomous spermatogenesis is likely adaptive. First, in some taxa it has been clearly demonstrated that both sperm morphs are required for successful fertilization, such as in the silkworm moth (*Bombyx mori*) [5]. Second, phylogenetic relationships indicate ancestral origins of dichotomous spermatogenesis and continued maintenance during evolution. For example, dichotomous spermatogenesis is present throughout Lepidoptera, with the sole exception of two species within the most basal suborder of this group. Although multiple independent origins of sperm heteromorphism in Lepidoptera has yet to be formally ruled out, a single ancestral origin is by far the most parsimonious explanation [6]. Third, the ratio or eupyrene to apyrene varies substantively across Lepidoptera but is relatively constant within species, including several cases where apyrene comprise up to 99% of the sperm produced [7]. While variation in the relative production of each sperm morph is not in itself incompatible with stochastic processes, such as drift, it is nearly impossible to reconcile the disproportionate investment in apyrene without acknowledging that they contribute in some fundamental way to reproductive fitness. Although far from definitive, it has also been suggested that this marked variability across species is consistent with ongoing diversifying selection [6]. Arriving at an understanding of apyrene function may be further complicated by the possibility that parasperm are generally more likely to acquire lineage specific functionalities [8].

To better understand the molecular basis of dichotomous spermatogenesis, we recently conducted a proteomic and genomic characterization of sperm in *Manduca sexta* (hereafter *Manduca*) [9]. An important component of our analysis was to determine the taxonomic distribution of sperm proteins, which revealed an unexpectedly high number of proteins that possess little or no homology to proteins outside of Lepidoptera. Although the genetic mechanisms responsible for this observation were not specifically explored, this pattern is consistent with genetic novelty associated with dichotomous spermatogenesis in Lepidoptera. Importantly, Lepidoptera specific sperm proteins were also determined to be significantly more abundant than other components of the *Manduca* sperm proteome, suggesting the function of some of these in apyrene sperm, which are by far the predominant morph in sperm samples in this species.

To provide a deeper understanding of the role of genetic novelty and genomic diversification in the evolution of dichotomous spermatogenesis, we have characterized the sperm proteome of the Monarch butterfly (*Danaus plexippus*; hereafter Monarch). In addition to its phylogenetic position and its continued development as a model butterfly species, we have pursued this species because of its distinct mating behavior. Unlike most other Lepidopteran species, male Monarch butterflies employ a strategy of coercive mating, resulting in female Monarchs to remate frequently [10]. In contrast, female remating is rare in *Manduca sexta* (hereafter *Manduca*) and, as in many other Lepidoptera, females attract males via attractants and calling behavior [11]. Interestingly, cessation of calling appears to be governed by molecular factors present in sperm or seminal fluid [12] and, as a consequence, non-virgin females rarely remate. Despite these behavioral differences, the proportion of eupyrene and apyrene produced is quite similar between these two species (~95-96%) [7,13]. Thus, our focus on Monarch is motivated both by their disparate, polyandrous mating system and their utility as a representative butterfly species for comparative analyses with *Manduca*. Therefore, the overarching aims of this study were to (**1**) characterize the sperm proteome of the Monarch butterfly and compare it with the previously characterized sperm proteome of *Manduca*, (**2**) contrast patterns of orthology across diverse insect genomes between the sperm proteome and remainder of genes in the genome and (**3**) analyze genome-wide homology to assess the contribution of evolutionary genetic novelty to Lepidopteran sperm composition.

## Methods

### Butterfly rearing and sperm purification

Adult male Monarch butterflies, kindly provided by MonarchWatch (Lawrence, Kansas), were dissected between 5 and 10 days post eclosion. The sperm contents of seminal vesicles, including both apyrene and eupyrene sperm, were dissected via a small incision in the mid to distal region of the seminal vesicle. Samples were rinsed in phosphate buffer solution and pelleted via centrifugation (2 minutes at 15,000 rpm) three times to produce a purified sperm sample. Sperm samples from 3-5 males were pooled to form three biological replicates [14].

### Protein Preparation and 1-Dimensional SDS Page

Samples were solubilized in 2X LDS sample buffer, as per manufacturers’ instructions (Invitrogen, Inc) before quantification via the EZA Protein Quantitation Kit (Invitrogen, Inc). Protein fluorescence was measured using a Typhoon Trio+ (Amersham Biosciences/GE Healthcare) with 488 nm excitation and a 610nm bandpass filter. Fluorescence data was analyzed using the ImageQuant TL software. Three replicates of 25ug of protein were separated on a 1 mm 10% NuPAGE Novex Bis-Tris Mini Gel set up using the XCell SureLock Mini-Cell system (Invitrogen) as per manufacturer instructions for reduced samples. Following electrophoresis, the gel was stained using SimplyBlue SafeStain (Invitrogen, Inc) and destained as per manufacturer instructions. Each lane on the resulting gel (containing sample form a single replicate) was sliced into four comparable slices, producing 12 gel fractions independent tandem mass spectrometry analysis.

### Tandem mass spectrometry (MS/MS)

Gel fractions were sliced into 1 mm^2^ pieces for in-gel trypsin digestion. Gel fractions were reduced (DDT) and alkylated (iodoacetamide) before overnight incubation with trypsin at 37 °C. All LC-MS/MS experiments were performed using a Dionex Ultimate 3000 RSLC nanoUPLC (Thermo Fisher Scientific Inc, Waltham, MA, USA) system and a QExactive Orbitrap mass spectrometer (Thermo Fisher Scientific Inc, Waltham, MA, USA). Separation of peptides was performed by reverse-phase chromatography at a flow rate of 300nL/min and a Thermo Scientific reverse-phase nano Easy-spray column (Thermo Scientific PepMap C18, 2μm particle size, 100A pore size, 75mm i.d. × 50cm length). Peptides were loaded onto a pre-column (Thermo Scientific PepMap 100 C18, 5μm particle size, 100A pore size, 300mm i.d. × 5mm length) from the Ultimate 3000 autosampler with 0.1% formic acid for 3 minutes at a flow rate of 10 μL/min. After this period, the column valve was switched to allow elution of peptides from the pre-column onto the analytical column. Solvent A was water plus 0.1% formic acid and solvent B was 80% acetonitrile, 20% water plus 0.1% formic acid. The linear gradient employed was 2-40% B in 30 minutes. The LC eluant was sprayed into the mass spectrometer by means of an Easy-spray source (Thermo Fisher Scientific Inc.). All m/z values of eluting ions were measured in an Orbitrap mass analyzer, set at a resolution of 70000. Data dependent scans (Top 20) were employed to automatically isolate and generate fragment ions by higher energy collisional dissociation (HCD) in the quadrupole mass analyzer and measurement of the resulting fragment ions was performed in the Orbitrap analyzer, set at a resolution of 17500. Peptide ions with charge states of 2+ and above were selected for fragmentation. The mass spectrometry proteomics data have been deposited to the ProteomeXchange Consortium via the PRIDE partner repository with the dataset identifier PXD006454 [15].

### MS/MS data analysis

MS/MS data was analyzed using X!Tandem and Comet algorithms within the Trans-Proteomic Pipeline (v 4.8.0) [16]. Spectra were matched against the *D. plexippus* official gene set 2 (OGS2) predicted protein set (downloaded from http://Monarchbase.umassmed.edu, last updated in 2012) with a fragment ion mass tolerance of 0.40 Da and a parent monoisotopic mass error of ±10 ppm. For both X!tandem and Comet, iodoacetamide derivative of cysteine was specified as a fixed modification, whereas oxidation of methionine was specified as a variable modification. Two missed cleavages were allowed and non-specific cleavages were excluded from the analysis. False Discovery Rates (FDRs) were estimated using a decoy database of randomized sequence for each protein in the annotated protein database. Peptide identifications were filtered using a greater than 95.0% probability based upon PeptideProphet [17] and the combined probability information from X!Tandem and Comet using Interprophet. Protein assignments were accepted if greater than 99.0%, as specified by the ProteinProphet [18] algorithms respectively. Proteins that contained identical peptides that could not be differentiated based on MS/MS analysis alone were grouped to satisfy the principles of parsimony. Protein inclusion in the proteome was based on the following stringent criteria: (**1**) identification in 2 or more biological replicates or (**2**) identification in a single replicate by 2 or more unique peptides. To identify post-translation modifications (PTMs) of proteins, X!Tandem and Comet were rerun allowing for variable phosphorylation of serine, threonine and tyrosine residues and acetylation of lysine residues. PTM locations were identified using PTMprophet in both the Monarch data presented here and a comparable dataset in *M. sexta* [19].

### APEX protein quantitation and analysis

Relative compositional protein abundance was quantified using the APEX Quantitative Proteomics Tool [20]. The training dataset was constructed using fifty proteins with the highest number of uncorrected spectral counts (*n_i_*), and identification probabilities. All 35 physicochemical properties available in the APEX tool were used to predict peptide detection/non-detection. Protein detection probabilities (O_i_) were computed using proteins with identification probabilities over 99% and the Random Forest classifier algorithm. APEX protein abundances were calculated using a merged protXML file generated by the ProteinProphet algorithm. The correlation in APEX abundance estimates of orthologous proteins in Monarch and *Manduca* (abundance estimates from Whittington et al. 2015) were normalized, log transformed and assessed using linear regression. Differential protein abundance was analyzed using corrected spectral counts and the R (v 3.0.0) package EdgeR [21]. Results were corrected for multiple testing using the Benjamini Hochberg method within EdgeR.

### Lift-over between D. plexippus version 1 and 2 gene sets

Two versions of gene models and corresponding proteins are currently available for *D. plexippus*. Official gene set one (OGS1) was generated using the genome assembly as initially published [22], while the more recent official gene set 2 (OGS2) was generated along with an updated genome assembly [23]. While our proteomic analysis employs the more recent OGS2 gene models, at the time of our analysis only OGS1 gene models were included in publicly available databases for gene function and orthology (e.g. Uniprot and OrthoDB). In order to make use of these public resources, we assigned OGS2 gene models to corresponding OGS1 gene models by sequence alignment. Specifically, OGS2 coding sequences (CDS) were aligned to OGS1 CDS using BLAT [24], requiring 95% identity; the best aligning OGS1 gene model was assigned as the match for the OGS2 query. In this way, we were able to link predictions of OGS1 gene function and orthology in public databases to OGS2 sequences in our analysis. Of the 584 OGS2 loci identified in the sperm proteome 18 could not be assigned to an OGS1 gene.

### Functional annotation and enrichment analysis

Two approaches were employed for functionally annotating *D. plexippus* sperm protein sequences. First, we obtained functional annotations assigned by Uniprot to corresponding *D. plexippus* OGS1 protein sequences (Additional file 1)[25]. Additionally we used the Blast2GO software to assign descriptions of gene function and also gene ontology categories [26]. The entire set of predicted protein sequences from OGS2 were BLASTed against the GenBank non-redundant protein database with results filtered for E<10^−5^, and also queried against the InterPro functional prediction pipeline [27]. Functional enrichment of GO terms present in the sperm proteome relative to the genomic background was performed using Blast2GO’s implementation of a Fisher’s exact test with a false discovery rate of 0.01%.

### Orthology predictions and analysis

Two approaches were employed for establishing orthology among proteins from different species. First, we used the proteinortho pipeline [28] to assess 3-way orthology beween *D. plexippus* OGS2, *M. sexta* OGS1 [29], and *D. melanogaster* (flybase r6.12) gene sets. Proteinortho uses a reciprocal blast approach to cluster genes with significant sequence similarities into clusters to identify orthologs and paralogs. For each species, genes with protein isoforms were represented by the longest sequence in the proteinortho analysis. *D. melanogaster* and *M. sexta* ortholog predictions were then cross referenced to the published sperm of these two species [9,30], allowing a three-way assessment of orthology in relation to presence in the sperm proteome. Using proteinortho allowed the direct analysis of the *D. plexippus* OGS2 sequences, which were not analyzed for homology in OrthoDB8 [31].

A taxonomically broader set of insect ortholog relationships was obtained from OrthoDB8 and used to assess the proportion of orthologs among sperm proteins relative to the genomic background. A randomized sampling procedure was used to determine the null expectation for the proportion of orthologous proteins found between *D. plexippus* and the queried species. A set of 584 proteins, the number equal to detected *D. plexippus* sperm proteins, was randomly sampled 5000 times from the entire Monarch OGS2 gene set. For each sample, the proportion of genes with an ortholog reported in OrthoDB8 was calculated, yielding a null distribution for the proportion of orthologs expected between *D. plexippus* and the queried species. For each query species, the observed proportion of orthologs in the sperm proteome was compared to this null distribution to determine whether the sperm proteome had a different proportion of orthologs than expected and to assign significance. Comparisons were made to 12 other insect species, reflecting five insect orders: Lepidoptera (*Heliconius melpomene, M. sexta, Plutella xylostella, Bombyx mori*), Diptera (*Drosophila melanogaster, Anopheles gambiae*), Hymenoptera (*Apis mellifera, Nasonia vitripennis*), Coleoptera (*Tribolium castaneum, Dendroctonus ponderosae*), and Hemiptera (*Acyrthosiphon pisum, Cimex lectularius*).

### Phylogenetic distribution and homology of sperm proteins

The taxonomic distribution of sperm proteins was determined by BLASTp analyses (statistical cut off of *e* < 10^−5^ and query coverage of ≥50%) against the protein data sets of the following taxa: butterflies (*Heliconius melpomene, Papilio xuthus, Lerema accius*), Lepidoptera (*M. sexta, Amyleios transitella, Plutella xylostella*), Diptera (*D. melanogaster*), Mecopterida (*Tribolium casteneum*), and Insecta (*Apis mellifera, Pediculus humanus, Acyrthosiphon pisum, Zootermopsis nevadensis*). Lepidopteran species were chosen to maximize species distribution across the full phylogenetic breadth of Lepidoptera, while also utilizing the most comprehensively annotated genomes based on published CEGMA scores (http://lepbase.org, [32]). Taxonomically restricted proteins were defined as those identified consistently across a given phylogenetic range without homology in any outgroup species. Proteins exhibiting discontinuous phylogenetic patterns of conservation were considered unresolved. Percentage identity information from BLAST searches between Monarch and *Manduca* were averaged for those proteins identified as Lepidoptera specific, those not specific to the Lepidoptera but with resolved taxonomic distribution, those with identified *Drosophila* orthology, and those without orthology in *Drosophila*. Mann-Whitney U tests were conducted to compare each average to the average percentage identity of Lepidoptera specific proteins. Bonferonni corrections were employed to cases of multiple testing.

## Results and Discussion

### Monarch sperm proteome

Characterization of the Monarch sperm proteome as part of this study, in conjunction with our previous analysis in *Manduca* [9], allowed us to conduct the first comparative analysis of sperm in Lepidoptera, and in insects more broadly, to begin to assess the origin and evolution of dichotomous spermatogenesis at the genomic level. Tandem mass spectrometry (MS/MS) analysis of Monarch sperm, purified in triplicate, identified 380 proteins in two or more replicates and 553 proteins identified by two or more unique peptides in a single replicate. Together this yielded a total of 584 high confidence protein identifications (Additional file 2). Of these, 41% were identified in all three biological replicates. Comparable with our previous analysis of *Manduca* sperm, proteins were identified by an average of 7.9 unique peptides and 21.1 peptide spectral matches. This new dataset thus provides the necessary foundation to refine our understanding of sperm composition at the molecular level in Lepidoptera. (Note: *Drosophila melanogaster* gene names will be used throughout the text where orthologous relationships exist with named genes; otherwise Monarch gene identification numbers will be used.)

### Gene Ontology analysis of molecular composition

Gene ontology (GO) analyses were first conducted to confirm the similarity in functional composition between the Monarch and other insect sperm proteomes. Biological process analyses revealed a significant enrichment for several metabolic processes, including the tricarboxylic acid (TCA) cycle (p= 2.22E-16), electron transport chain (p= 9.85E-18), oxidation of organic compounds (p= 1.33E-25) and generation of precursor metabolites and energy (p= 1.09E-30) (Fig. 1a). GO categories related to the TCA cycle and electron transport have also been identified to be enriched in the *Drosophila* and *Manduca* sperm proteomes [9]. Generation of precursor metabolites and energy, and oxidation of organic compounds are also the two most significant enriched GO terms in the *Drosophila* sperm proteome [30]. Thus, broad metabolic functional similarities exist between the well-characterized insect sperm proteomes.

**Figure 1.**
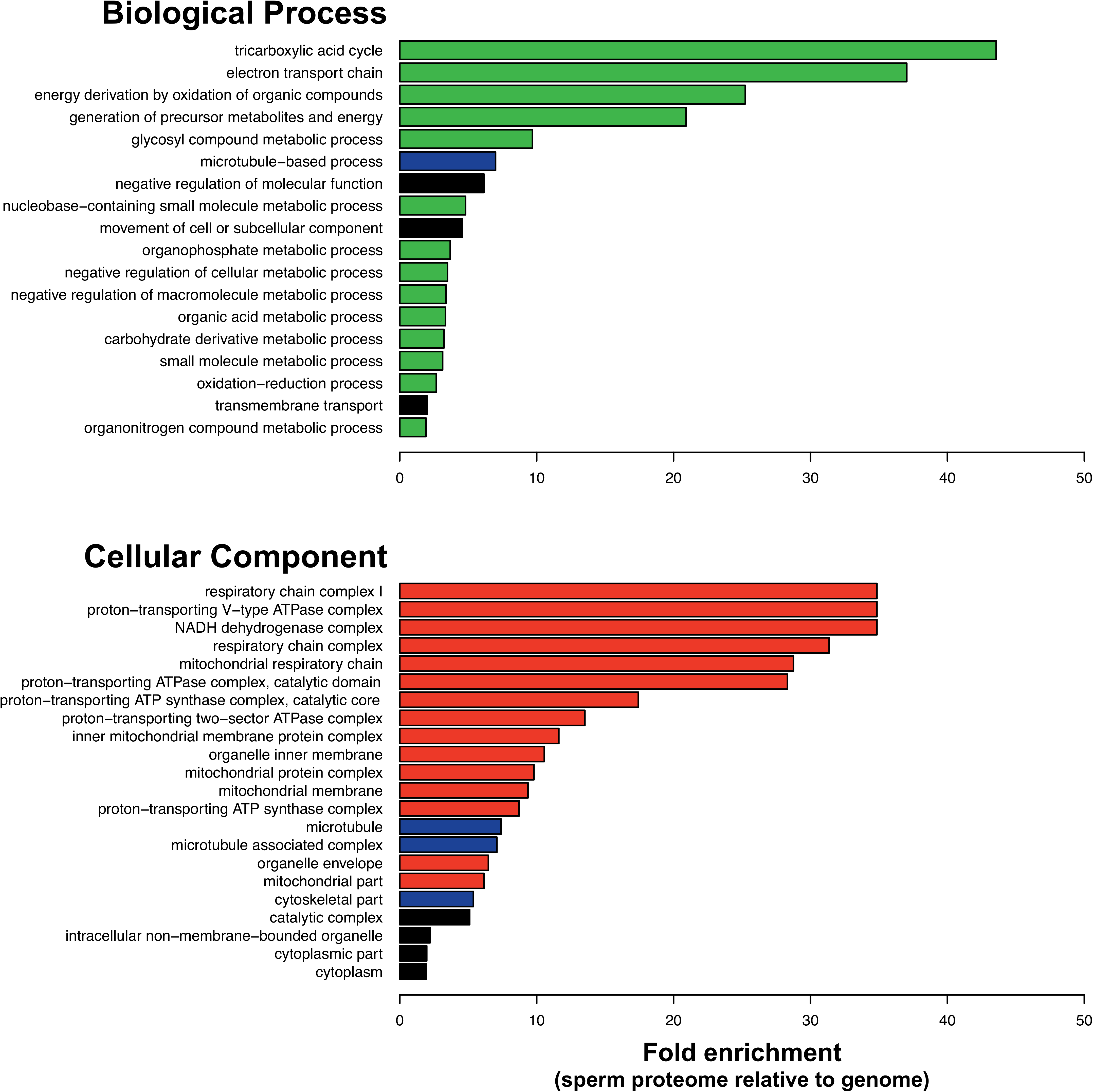
Functional enrichment within the Monarch sperm proteome. Biological Process and Cellular Component Gene Ontology enrichments in the sperm proteome relative to the whole genome were conducted using Blast2GO’s Fisher’s exact test with a false discovery rate of 0.01%. All categories displayed achieved significance. Functional categories directly relevant to sperm biology are indicated: metabolism (green), structural (blue) and mitochondria associated components (red).

An enrichment of proteins involved in microtubule-based processes was also observed, a finding that is also consistent with previously characterized insect sperm proteomes. Amongst the proteins identified are cut up (ctp), a dynein light chain required for spermatogenesis [33], actin 5 (Act5), which is involved in sperm individualization [34], and DPOGS212342, a member of the recently expanded X-linked *tektin* gene family in *Drosophila* sperm [35]. Although functional annotations are limited amongst the 10% most abundant proteins (see below), several contribute to energetic and metabolic pathways. For example, stress-sensitive B (sesB) and adenine nucleotide translocase 2 (Ant2) are gene duplicates that have been identified in the *Drosophila* sperm proteome and, in the case of Ant2, function specifically in in mitochondria during spermatogenesis [36]. Also identified was Bellwether (blw), an ATP synthetase alpha chain which is required for spermatid development [37].

The widespread representation of proteins functioning in mitochondrial energetic pathways is consistent with the contribution of giant, fused mitochondria (i.e. nebenkern) in flagellum development and presence of mitochondrial derivatives in mature spermatozoa (Fig 1a-b) [38]. During lepidopteran spermatogenesis, the nebenkern divides to form two derivatives, which flank the axoneme during elongation; ultrastructure and size of these derivatives varies greatly between species and between the two sperm morphs [7]. In *Drosophila*, the nebenkern acts as both an organizing center for microtubule polymerization and a source of ATP for axoneme elongation, however it is unclear to what extent these structures contribute to energy required for sperm motility. Of particular note is the identification of porin, a voltage-gated anion channel that localizes to the nebenkern and is critical for sperm mitochondrion organization and individualization [39]. Consistent with these patterns, Cellular Component analysis also revealed a significant enrichment of proteins in a broad set of mitochondrial structures and components, including the respiratory chain complex I (p = 7.73E-09), proton-transporting V-type ATPase complex (p = 9.90E-08) and the NADH dehydrogenase complex (p = 7.73E-09) (Fig. 1b). Aside from those categories relating to mitochondria, a significant enrichment was also observed amongst categories relating to flagellum structure, including microtubule (p = 5.43E-18) and cytoskeleton part (p = 2.54E-12). The two most abundant proteins in the proteome identified in both Monarch and *Manduca*, beta tubulin 60D (βTub60D) and alpha tubulin 84B (βTub84B), contributed to these GO categories. αTub84B is of particular interest as it performs microtubule functions in the post-mitotic spermatocyte, including the formation of the meiotic spindle and sperm tail elongation [40].

Molecular Function GO analysis revealed an enrichment of oxidoreductase proteins acting on NAD(P)H (p = 7.06E-19), as well as more moderate enrichments in several categories relating to peptidase activity or regulation of peptidase activity (data not shown). The broad representation of proteins involved in proteolytic activity is worthy of discussion, not solely because these classes of proteins are abundant in other sperm proteomes, but also because proteases are involved in the breakdown of the fibrous sheath surrounding Lepidoptera eupyrene sperm upon transfer to the female [7]. This process has been attributed to a specific ejaculatory duct trypsin-like arginine C-endopeptidase (initiatorin) in the silkworm (*B. mori*) [41] and a similar enzymatic reaction is needed for sperm activation in *Manduca* [42]. Blast2GO analyses identified three serine-type proteases in the top 5% of proteins based on abundance, including a chymotrypsin peptidase (DPOGS213461) and a trypsin precursor (DPOGS205340). These highly abundant proteases, particularly those that were also identified in *Manduca* (two of the most abundant proteases and 10 in total), are excellent candidates for a sperm activating factor(s) in Lepidoptera.

### Conservation of Lepidoptera Sperm Proteomes

Our previous analysis of *Manduca* was the first foray into the molecular biology of Lepidopteran sperm and was motivated by our interest in the intriguing heteromorphic sperm system that is found in nearly all species in this order [7]. Here we have aimed to delineate the common molecular components of lepidopteran sperm through comparative analyses. Orthology predictions between the two species identified relationships for 405 (69%) Monarch sperm proteins and 298 of these (73.5%) were previously identified by MS/MS in the *Manduca* sperm proteome [9]. An identical analysis in *Drosophila* identified 203 (35%) Monarch proteins with orthology relationships, including 107 (52.7%) that were previously characterized as components of the *Drosophila* sperm proteome [30,43]. Thus, and as would be expected given the taxonomic relationship of these species, there is a significantly greater overlap in sperm components between the two Lepidopteran species (two tailed Chi-square = 25.55, d.f. = 1, p < 0.001).

Recent comparative analyses of sperm composition across mammalian orders successfully identified a conserved “core” sperm proteome comprised of more slowly evolving proteins, including a variety of essential structural and metabolic components [61]. To characterize the “core” proteome in insects, we conducted a GO analysis using *Drosophila* orthology, ontology and enrichment data to assess the molecular functionality of proteins identified in the proteome of all three insect species. This revealed a significant enrichment for proteins involved in cellular respiration (p= 4.41e-21), categories associated with energy metabolism, including ATP metabolic process (p= 1.64e-15), generation of precursor metabolites and energy (p= 9.77e-21), and multiple nucleoside and ribonucleoside metabolic processes. Analysis of cellular component GO terms revealed a significant enrichment for mitochondrion related proteins (p= 3.72e-22), respiratory chain complexes (p= 8.25e-12), dynein complexes (p= 1.37e-5), and axoneme (p=3.31e-6). These GO category enrichments are consistent with a core set of metabolic, energetic, and structural proteins required for general sperm function. Similar sets core sperm proteins have been identified in previous sperm proteome comparisons [9,30,43,44]. Among this conserved set are several with established reproductive phenotypes in *Drosophila*. This includes proteins associated with sperm individualization, including cullin3 (Cul3) and SKP1-related A (SkpA), which acts in cullin-dependent E3 ubiquitin ligase complex required for caspase activity in sperm individualization [45], gudu, an Armadillo repeat containing protein [46], and porin (mentioned previously) [39]. Two proteins involved in sperm motility were also identified: dynein axonemal heavy chain 3 (dnah3) [47] and an associated microtubule-binding protein growth arrest specific protein 8 (Gas8) [48].

### Comparative analysis of protein abundance

Despite the more proximate link between proteome composition and molecular phenotypes, transcriptomic analyses far outnumber similar research using proteomic approaches. Nonetheless, recent work confirms the utility of comparative evolutionary proteomic studies in identifying both conserved [49] and diversifying proteomic characteristics [50]. We have previously demonstrated a significant correlation in protein abundance between *Manduca* and *Drosophila* sperm, although this analysis was limited by the extent of orthology between these taxa [9]. To further investigate the evolutionary conservation of protein abundance in sperm, a comparison of normalized abundance estimates between Monarch and *Manduca* revealed a highly significant correlation (R^2^= 0.43, p=<1x10^−15^) (Fig. 2A). We note that this correlation is based on semi-quantitative estimates [20] and would most likely be stronger if more refined absolute quantitative data were available. Several proteins identified as highly abundant in both species are worthy of further mention. *Sperm leucyl aminopeptidase 7* (*S-Lap7*) is a member of gene family first characterized in *Drosophila* that has recently undergone a dramatic expansion, is testis-specific in expression and encodes the most abundant proteins in the *D. melanogaster* sperm proteome [51]. As would be expected, several microtubule structural components were also amongst the most abundant proteins (top 20), including αTub84B and tubulin beta 4b chain-like protein, as well as succinate dehydrogenase subunits A and B (SdhA and SdhB), porin, and DPOGS202417, a trypsin precursor that undergoes conserved post translational modification (see below).

**Figure 2.**
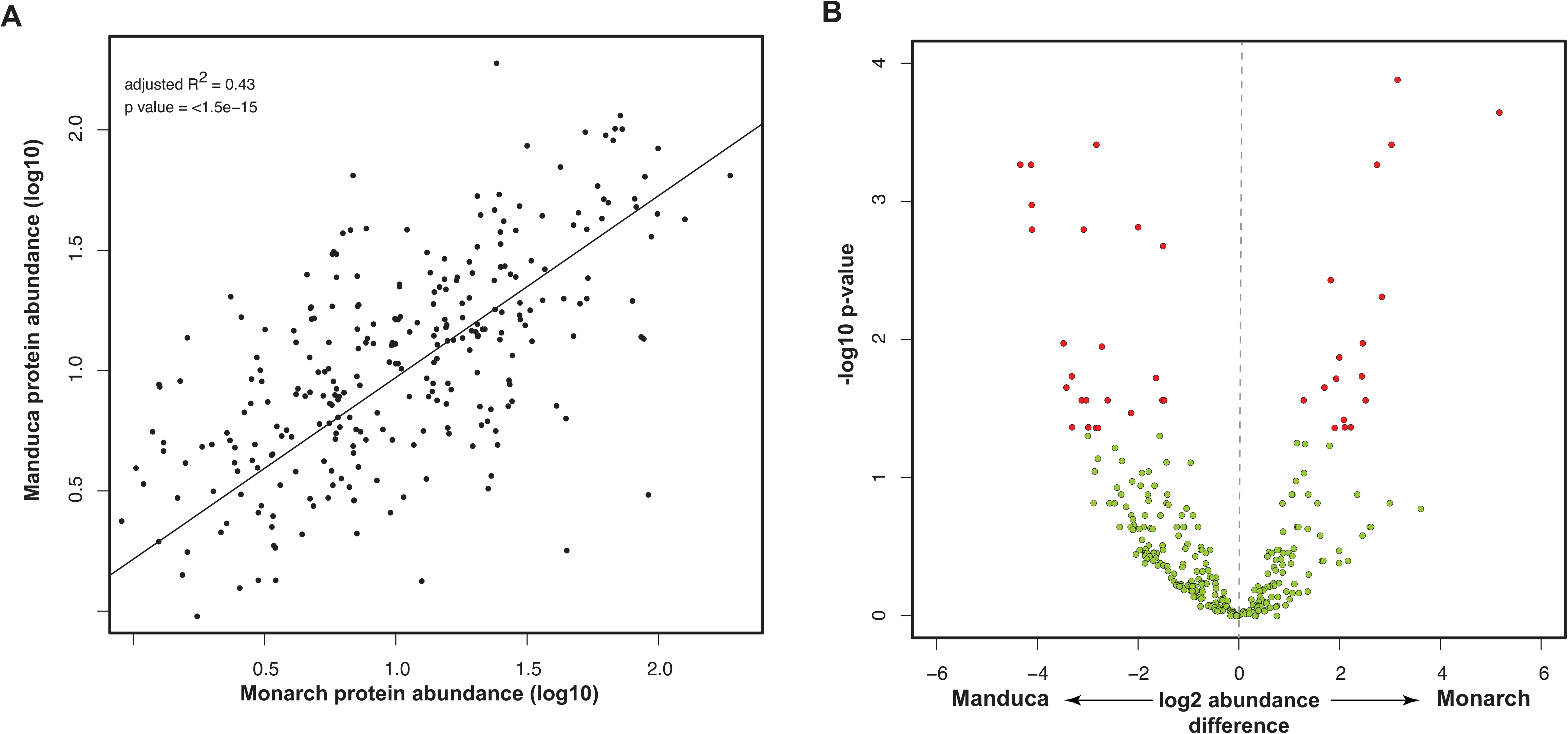
Conservation of Monarch and *Manduca* sperm composition. (A) Linear regression analysis of protein abundance estimates for proteins identified in both species reveals a significant correlation. (B) Differential abundance analysis using EdgeR revealed 45 significant proteins after Benjamini Hochberg multiple testing correction. Proteins significantly different in abundance between species are shown in red, nonsignificant proteins are shown in green. Proteins with negative values are more abundant in *Manduca* whereas positive values are more abundant in Monarch.

We next sought to identify proteins exhibiting differential abundance between the two species. As discussed earlier, Monarch and *Manduca* have distinct mating systems; female Monarch butterflies remate considerably more frequently than *Manduca* females, increasing the potential for sperm competition [10]. These differences may be reflected in molecular diversification in sperm composition between species. An analysis of differential protein abundance identified 45 proteins with significant differences (P<0.05; Fig. 2B), representing 7% of the proteins shared between species (Additional file 3). No directional bias was observed in the number of differentially abundant proteins (one-tail Binomial test; p value = 0.2757). Several of these proteins are worthy of further discussion given their role in sperm development, function or competitive ability. Proteins identified as more abundant in the Monarch sperm proteome were heavily dominated by mitochondrial NADH dehydrogenase subunits (subunits ND-23, ND-24, ND-39, and ND-51) and other mitochondria-related proteins, including ubiquinol-cytochrome c reductase core protein 2 (UQCR-C2), cytochrome C1 (Cyt-C1), and glutamate oxaloacetate transaminase 2 (Got2). Additionally, two proteins with established sperm phenotypes were identified as more abundant in *Manduca*. These included dynein light chain 90F (Dlc90F), which is required for proper nuclear localization and attachment during sperm differentiation [52], and cut up (ctp), a dynein complex subunit involved in nucleus elongation during spermiogenesis [33]. Serine protease immune response integrator (spirit) is also of interest considering the proposed role of endopeptidases in Lepidoptera sperm activation [41,42]. Although it would be premature to draw any specific conclusions, some of these proteins play important mechanistic roles in sperm development and function and will be of interest for more targeted functional studies.

### Post-translational modification of sperm proteins

During spermatogenesis, the genome is repackaged and condensed on protamines and the cellular machinery required for protein synthesis are expelled. Consequently, mature sperm cells are considered primarily quiescent [53]. Nonetheless, sperm undergo dynamic molecular transformations after they leave the testis and during their passage through the male and female reproductive tract [54]. One mechanism by which these modifications occur is via post translational modification (PTM), which can play an integral part in the activation of sperm motility and fertilization capacity [55,56]. Analysis of PTMs in Monarch identified 438 acetylated peptides within 133 proteins. Most notable among these are microtubule proteins, including alpha tubulin 84B (alphaTub84B), beta tubulin 60D (betaTub60D) and dyneins kl-3 and kl-5. Tubulin is a well-known substrate for acetylation, including the highly-conserved acetylation of N-terminus Lysine 40 of alphaTub84B. This modification is essential for normal sperm development, morphology and motility in mice [57]. A similar analysis in *Manduca* identified 111 acetylated peptides within 63 proteins. We found evidence for conserved PTMs within Lepidoptera in 19 proteins (36% of those identified in Monarch), including Lys40 of alphaTub84B.

In contrast to acetylation, only 75 Monarch sperm proteins showed evidence of phosphorylation, 53 of which were also modified in *Manduca* (36%). This included the ortholog of the Y-linked *Drosophila* gene WDY. Although a specific function for WDY in spermatogenesis has yet to be determined, WDY is expressed in a testis-specific manner and under positive selection in the *D. melanogaster* group [58]. The relative paucity of phosphorylation PTMs may reflect the fact that phosphorylation is one of the more difficult PTMs to identify with certainty via mass spectrometry based proteomics [59]. However, it is also noteworthy that sperm samples in this study were purified from the male seminal vesicle, and thus, before transfer to the female reproductive tract. Although far less is known about the existence of capacitation-like processes in insects, dynamic changes in the mammalian sperm phosphoproteome are associated with sperm capacitation and analogous biochemical alterations might occur within the female reproductive tract of insects [56]. We note that a similar extent of protein phosphorylation has been detected from *Drosophila* sperm samples purified in a similar manner (unpublished data; Whittington and Dorus). Lastly, identical acetylation and phosphorylation PTM patterns were identified for Monarch and *Manduca* HACP012 (DPOGS213379), a putative seminal fluid protein of unknown function previously identified in the Postman butterfly (*Heliconius melpomene*) [60,61]. The identification of HACP012 in sperm, in the absence of other seminal fluid components, is unexpected but its identification was unambiguous as it was amongst the most abundant 10% of identified Monarch proteins. Seminal protein HACP020 (DPOGS203866), which exhibits signatures of recent adaptive evolution [61], was also identified as highly abundant (5^th^ percentile overall); this suggests that some seminal fluid proteins may also be co-expressed in the testis and establish an association with sperm during spermatogenesis.

### Rapid evolution of genetic architecture

Rapid gene evolution [62] and gene creation/loss [63], including *de novo* gene creation [64], are predominant processes that contribute to the diversification of male reproductive systems. Our previous study identified an enrichment in the number of Lepidoptera specific proteins (*i.e*. those without homology outside of Lepidoptera) in the sperm proteome relative to other reproductive proteins. We were unable, however, to determine from a single species whether novel genes contributed to sperm biology more broadly across all Lepidoptera. Here we employed two comparative genomic approaches to confirm and expand upon our original observation. First, we obtained whole-genome orthology relationships between Monarch and nine species, representing five insect orders, and compared the proportion of the sperm proteome with orthologs to the whole genome using a random subsampling approach. No significant differences were observed for three of the four Lepidoptera species analyzed and an excess of orthology amongst sperm proteins was identified in the Postman butterfly (p < 0.05; Fig. 3). In contrast, we identified a significant deficit of sperm orthologs in all comparisons with non-Lepidopteran genomes (all p < 0.01). Orthology relationships in OrthoDB are established by a multi-step procedure involving reciprocal best match relationships between species and identity within species to account for gene duplication events since the last common ancestor. As such, the underrepresentation of orthology relationships is unlikely to be accounted for by lineage-specific gene duplication. Therefore, rapid evolution of sperm genes appears to be the most reasonable explanation for the breakdown of reciprocal relationships (see below). This conclusion is consistent with a diverse body of evidence that supports the influence of positive selection on male reproductive genes [62,65], including those functioning in sperm [43,66–68]. We note that we cannot rule out the influence of *de novo* creation but it is currently difficult to assess the contribution of this mechanism to the overall pattern.

**Figure 3.**
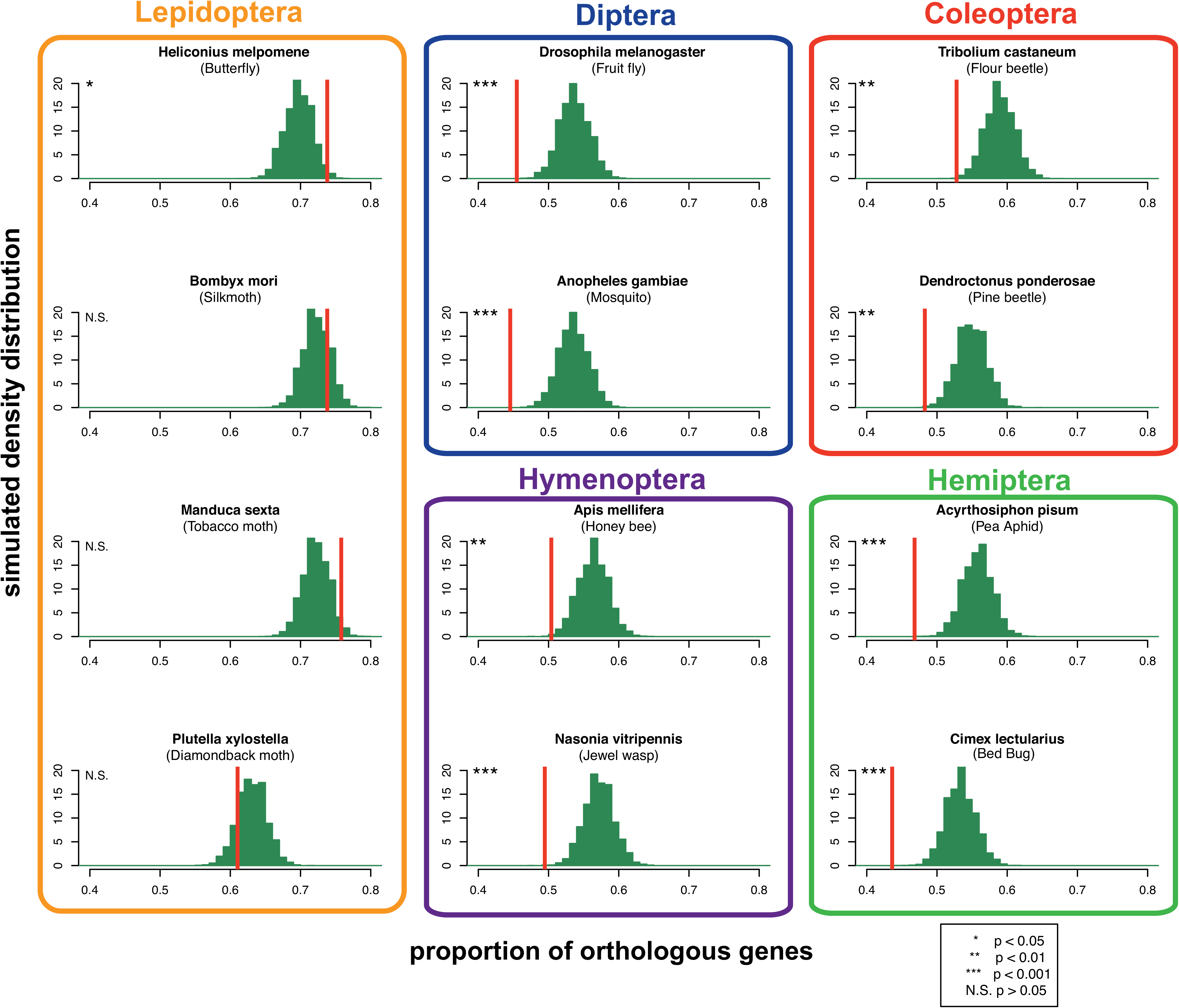
Sperm proteome orthology relationships in insects. Orthology relationships were curated from OrthoDB8 in 12 species, reflecting five insect orders: Lepidoptera, Diptera, Hymenoptera, Coleoptera, and Hemiptera. The distribution of expected orthology relationships for each species was determined by 5,000 randomized subsamples of Monarch genes not identified in the sperm proteome (green bars). The observed proportion of orthologs for the Monarch proteome are indicated (red line).

The second analysis aimed to characterize the distribution of taxonomically restricted Monarch sperm proteins using BLAST searches across 12 insect species. Based on the analysis above, our *a priori* expectation was that a substantial number of proteins with identifiable homology amongst Lepidoptera would be absent from more divergent insect species. This analysis identified a total of 45 proteins unique to Monarch, 140 proteins (23.9% of the sperm proteome) with no homology to proteins in non-Lepidopteran insect taxa and 173 proteins conserved across all species surveyed (Fig. 4a). Proteins with discontinuous taxonomic matches (n = 171) were considered “unresolved”. Although the number of Monarch-specific proteins is considerably higher than the eight *Manduca-specific* proteins found in our previous study, the number of Lepidoptera specific is comparable to our previous estimate in *Manduca* (n = 126). These observations support the hypothesis that a substantial subset of lepidopteran sperm proteins are likely to be rapidly evolving and thus exhibit little detectable similarity. To pursue this possibility, we estimated Lepidoptera specific protein divergence between Monarch and *Manduca* and compared the distribution of amino acid divergence to those proteins identified in other insect species (Fig. 4B). The average percentage identity of Lepidoptera specific proteins (55.1% ± 17.6) was significantly lower than all non-Lepidopteran specific proteins (74.6% ± 13.4, W=3074.5, p=3.35e-16), those with *Drosophila* orthology (75.5% ± 13.6, W=8285.5, p=<1x10^−15^), and those non-Lepidopteran specific proteins without *Drosophila* orthology (62.4% ± 18.4, W=2980, p=1.16e-5). Therefore, we can conclude that Lepidoptera specific proteins evolve more rapidly than other sperm proteins and that proteins with resolved orthology relationships in *Drosophila* experience higher levels of conservation than those that do not. To assess their potential contribution to sperm function, we used protein abundance as a general proxy in the absence of functional annotation for nearly all of these proteins. As was observed in Whittington *et al* [9], Lepidopteran specific proteins were found to be significantly more abundant than the remainder of the sperm proteome (D=0.2, p=0.0009, Fig. 4c).

**Figure 4.**
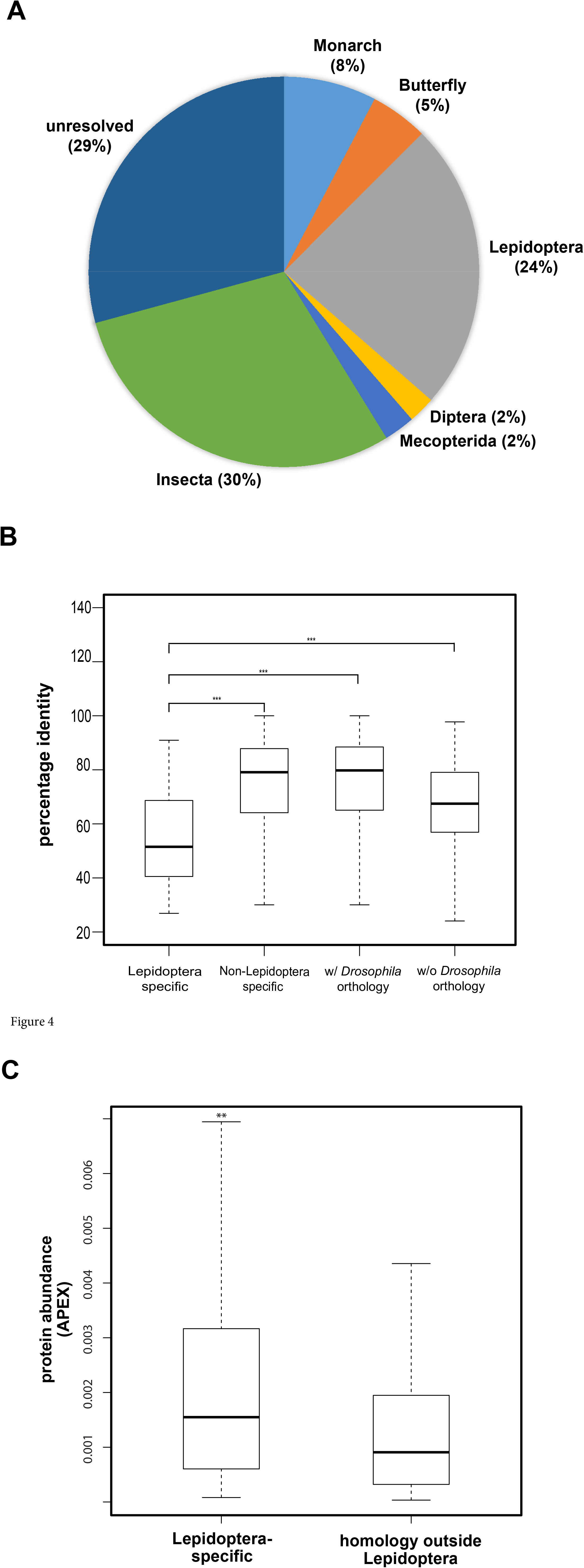
Taxonomic distribution of Monarch sperm protein homology in insects. (A) Pie chart displaying the taxonomical distribution of proteins homologous to the Monarch sperm proteome and those unique to Monarch. BLAST searches were conducted beginning with closely related butterfly species and sequentially through more divergent species in Mercopterida, Diptera and Insecta. In order to be considered Lepidoptera specific, a protein was required to be present in at least at least one butterfly other than Monarch and at least one moth species. Proteins with discontinuous taxonomic patterns of homology are included in the category “unresolved”. (B) Box plot showing percentage identity for BLAST hits between Monarch and *Manduca*. Percentage identities for proteins identified as specific to Lepidoptera, non-specific but with resolved taxonomic distribution, with identified orthology in *Drosophila*, and without orthology in *Drosophila* are shown. (C) Box plot displaying the distribution of protein abundance estimates for proteins present only in Lepidoptera and those with homology in other insects.

### Conclusion

This comparative proteomic analysis of heteromorphic sperm, a first of its kind, provides important perspective and insights regarding the functional and evolutionary significance of this enigmatic reproductive phenotype. Our analyses indicate that a substantial number of novel sperm genes are shared amongst Lepidoptera, thus distinguishing them from other insect species without dichotomous spermatogenesis, and suggest they are associated with heteromorphic spermatogenesis and the diversification of apyrene and eupyrene sperm. Our comparative and quantitative analyses, based on protein abundance measurements in both species, further suggests that some of these proteins contribute to apyrene sperm function and evolution. Given that apyrene sperm constitute the vast majority of cells in our co-mixed samples, it is reasonable to speculate that higher abundance proteins are either present in both sperm morphs or specific to apyrene cells and thus good candidates for further study in relation to apyrene sperm functionality.

## Declarations

Mass spectrometry data is publicly available through the ProteomeXchange Consortium (http://proteomecentral.proteomexchange.org) with the dataset identifier PXD006454. There are no financial or non-financial interests associated with this study. Funding for this study included Syracuse University support to SD, University of Kansas support to JW and a Syracuse University and Marilyn Kerr Fellowships to EW. We thank Monarch Watch and Channing Shives for support in rearing Monarch butterflies and Sheri Skerget for expert technical assistance. Computing for this project was performed on the Syracuse University Crush Virtual Research Cloud and the Community Cluster at the Center for Research Computing at the University of Kansas. TLK, JW and SD designed the study; TLK and JW purified samples for MS analysis; EW, DH, JW and SD analyzed the data; ECW, TLK, JW and SD wrote the manuscript.

## References

1. Pitnick S, Birkhead TR, Hosken DJ, editors. Sperm biology: an evolutionary perspective. 1st ed. Amsterdam: Academic Press/Elsevier; 2009.

2. Till-Bottraud I, Joly D, Lachaise D, Snook RR. Pollen and sperm heteromorphism: convergence across kingdoms? J. Evol. Biol. 2005;18:1–18.

3. Swallow JG, Wilkinson GS. The long and short of sperm polymorphisms in insects. Biol. Rev. Camb. Philos. Soc. 2002;77:153–82.

4. Ueber, M. F. Oligopyrene und apyrene spermien und über ihre entstehung, nach Beobachtungen an Paludina und Pygaera. Arch. Für Mikrosk. Anat. 1902;61:1–84.

5. Sahara K, Kawamura N. Double copulation of a female with sterile diploid and polyploid males recovers fertility in *Bombyx mori*. Zygote Camb. Engl. 2002;10:23–9.

6. Friedländer M. Control of the eupyrene–apyrene sperm dimorphism in *Lepidoptera*. J. Insect Physiol. 1997;43:1085–92.

7. Friedländer M, Seth RK, Reynolds SE. Eupyrene and apyrene sperm: Dichotomous spermatogenesis in *Lepidoptera*. Adv. Insect Physiol. 2005;32:206–308.

8. Snook RR, Hosken DJ, Karr TL. The biology and evolution of polyspermy: insights from cellular and functional studies of sperm and centrosomal behavior in the fertilized egg. Reproduction. 2011;142:779–92.

9. Whittington E, Zhao Q, Borziak K, Walters JR, Dorus S. Characterisation of the *Manduca sexta* sperm proteome: Genetic novelty underlying sperm composition in *Lepidoptera*. Insect Biochem. Mol. Biol. 2015;62:183–93.

10. Oberhauser K, Frey D. Coercive mating by overwintering male monarch butterflies. 1997 North Am. Conf. Monarch Butterfly. 1997. p. 67.

11. Sasaki M, Riddiford LM. Regulation of reproductive behaviour and egg maturation in the tobacco hawk moth, *Manduca sexta*. Physiol. Entomol. 1984;9:315–27.

12. Stringer IAN, Giebultowicz JM, Riddiford LM. Role of the bursa copulatrix in egg maturation and reproductive behavior of the tobacco hawk moth, *Manduca sexta*. Int. J. Invertebr. Reprod. Dev. 1985;8:83–91.

13. Solensky MJ, Oberhauser KS. Male monarch butterflies, *Danaus plexippus*, adjust ejaculates in response to intensity of sperm competition. Anim. Behav. 2009;77:465–72.

14. Karr TL, Walters JR. Panning for sperm gold: Isolation and purification of apyrene and eupyrene sperm from lepidopterans. Insect Biochem. Mol. Biol. 2015;63:152–8.

15. Vizcaíno JA, Csordas A, del-Toro N, Dianes JA, Griss J, Lavidas I, et al. 2016 update of the PRIDE database and its related tools. Nucleic Acids Res. 2016;44:11033–11033.

16. Deutsch EW, Mendoza L, Shteynberg D, Farrah T, Lam H, Tasman N, et al. A guided tour of the Trans-Proteomic Pipeline. Proteomics. 2010;10:1150–9.

17. Keller A, Nesvizhskii AI, Kolker E, Aebersold R. Empirical statistical model to estimate the accuracy of peptide identifications made by MS/MS and database search. Anal. Chem. 2002;74:5383–92.

18. Nesvizhskii AI, Keller A, Kolker E, Aebersold R. A statistical model for identifying proteins by tandem mass spectrometry. Anal. Chem. 2003;75:4646–58.

19. Shteynberg DD., Mendoza L, Slagel J, Lam H, Nesvizhskii AI, Moritz R. PTMProphet: TPP software for validation of modified site locations on post-translationally modified peptides. 60th American Society for Mass Spectrometry (ASMS) Annual Conference, Vancouver, Canada, 2012.

20. Braisted JC, Kuntumalla S, Vogel C, Marcotte EM, Rodrigues AR, Wang R, et al. The APEX quantitative proteomics tool: Generating protein quantitation estimates from LC-MS/MS proteomics results. BMC Bioinformatics. 2008;9:529.

21. Robinson MD, McCarthy DJ, Smyth GK. edgeR: a Bioconductor package for differential expression analysis of digital gene expression data. Bioinformatics. 2010;26:139–40.

22. Zhan S, Merlin C, Boore JL, Reppert SM. The Monarch butterfly genome yields insights into long-distance migration. Cell. 2011;147:1171–85.

23. Zhan S, Reppert SM. MonarchBase: the Monarch butterfly genome database. Nucleic Acids Res. 2013;41:D758–63.

24. Kent WJ. BLAT---The BLAST-Like alignment tool. Genome Res. 2002;12:656–64.

25. The UniProt Consortium. UniProt: a hub for protein information. Nucleic Acids Res. 2015;43:D204–12.

26. Conesa A, Gotz S, Garcia-Gomez JM, Terol J, Talon M, Robles M. Blast2GO: a universal tool for annotation, visualization and analysis in functional genomics research. Bioinformatics. 2005;21:3674–6.

27. Zdobnov EM, Apweiler R. InterProScan - an integration platform for the signature-recognition methods in InterPro. Bioinforma. Oxf. Engl. 2001;17:847–8.

28. Lechner M, Findeiß S, Steiner L, Marz M, Stadler PF, Prohaska SJ. Proteinortho: Detection of (co-)orthologs in large-scale analysis. BMC Bioinformatics. 2011;12:124.

29. Kanost MR, Arrese EL, Cao X, Chen Y-R, Chellapilla S, Goldsmith MR, et al. Multifaceted biological insights from a draft genome sequence of the tobacco hornworm moth, *Manduca sexta*. Insect Biochem. Mol. Biol. 2016;76:118–47.

30. Wasbrough ER, Dorus S, Hester S, Howard-Murkin J, Lilley K, Wilkin E, et al. The *Drosophila melanogaster* sperm proteome-II (DmSP-II). J. Proteomics. 2010;73:2171–85.

31. Waterhouse RM, Tegenfeldt F, Li J, Zdobnov EM, Kriventseva EV. OrthoDB: a hierarchical catalog of animal, fungal and bacterial orthologs. Nucleic Acids Res. 2013;41:D358–65.

32. Challis RJ, Kumar S, Dasmahapatra KKK, Jiggins CD, Blaxter M. Lepbase: The Lepidopteran genome database. bioRxiv. 2016; doi: 10.1101/056994

33. Joti P, Ghosh-Roy A, Ray K. Dynein light chain 1 functions in somatic cyst cells regulate spermatogonial divisions in *Drosophila*. Sci. Rep. 2011;1:173.

34. Noguchi T. A role for actin dynamics in individualization during spermatogenesis in *Drosophila melanogaster*. Development. 2003;130:1805–16.

35. Dorus S, Freeman ZN, Parker ER, Heath BD, Karr TL. Recent origins of sperm genes in Drosophila. Mol. Biol. Evol. 2008;25:2157–66.

36. Terhzaz S, Cabrero P, Chintapalli VR, Davies S-A, Dow JAT. Mislocalization of mitochondria and compromised renal function and oxidative stress resistance in *Drosophila* SesB mutants. Physiol. Genomics. 2010;41:33–41.

37. Castrillon DH, Gönczy P, Alexander S, Rawson R, Eberhart CG, Viswanathan S, et al. Toward a molecular genetic analysis of spermatogenesis in *Drosophila melanogaster:* characterization of male-sterile mutants generated by single P element mutagenesis. Genetics. 1993;135:489–505.

38. Tokuyasu KT. Dynamics of spermiogenesis in *Drosophila melanogaster*. VI. Significance of “onion” nebenkern formation. J. Ultrastruct. Res. 1975;53:93–112.

39. Park J, Kim Y, Choi S, Koh H, Lee S-H, Kim J-M, et al. *Drosophila* Porin/VDAC affects mitochondrial morphology. PloS One. 2010;5:e13151.

40. Hutchens JA, Hoyle HD, Turner FR, Raff EC. Structurally similar *Drosophila* alpha-tubulins are functionally distinct in vivo. Mol. Biol. Cell. 1997;8:481–500.

41. Osanai M, Kasuga H, Aigaki T. Induction of motility of apyrene spermatozoa and dissociation of Eupyrene sperm bundles of the silkmoth, *Bombyx mori*, by initiatorin and trypsin. Invertebr. Reprod. Dev. 1989;15:97–103.

42. Friedländer M, Jeshtadi A, Reynolds SE. The structural mechanism of trypsin-induced intrinsic motility in *Manduca sexta* spermatozoa in vitro. J. Insect Physiol. 2001;47:245–55.

43. Dorus S, Busby SA, Gerike U, Shabanowitz J, Hunt DF, Karr TL. Genomic and functional evolution of the *Drosophila melanogaster* sperm proteome. Nat. Genet. 2006;38:1440–5.

44. Rettie EC, Dorus S. *Drosophila* sperm proteome evolution: Insights from comparative genomic approaches. Spermatogenesis. 2012;2:213–23.

45. Arama E, Bader M, Rieckhof GE, Steller H. A Ubiquitin Ligase Complex Regulates Caspase Activation During Sperm Differentiation in *Drosophila*. Bach E, editor. PLoS Biol. 2007;5:e251.

46. Cheng W, Ip YT, and Xu Z. Gudu, an Armadillo repeat-containing protein, is required for spermatogenesis in *Drosophila*. Gene. 2013;531:294–300.

47. Karak S, Jacobs JS, Kittelmann M, Spalthoff C, Katana R, Sivan-Loukianova E, et al. Diverse roles of axonemal dyneins in *Drosophila* auditory neuron function and mechanical amplification in hearing. Sci. Rep. 2015;5:17085.

48. Yeh S-D, Chen Y-J, Chang ACY, Ray R, She B-R, Lee W-S, et al. Isolation and properties of *Gas8*, a growth arrest-specific gene regulated during male gametogenesis to produce a protein associated with the sperm otility apparatus. J. Biol. Chem. 2002;277:6311–7.

49. Bayram HL, Claydon AJ, Brownridge PJ, Hurst JL, Mileham A, Stockley P, et al. Cross-species proteomics in analysis of mammalian sperm proteins. J. Proteomics. 2016;135:38–50.

50. Vicens A, Borziak K, Karr TL, Roldan ERS, Dorus S. Comparative sperm proteomics in mouse species with divergent mating systems. Mol. Biol. Evol. 2017; doi: 10.1093/molbev/msx084

51. Dorus S, Wilkin EC, Karr TL. Expansion and functional diversification of a leucyl aminopeptidase family that encodes the major protein constituents of *Drosophila* sperm. BMC Genomics. 2011;12.

52. Li M, Serr M, Newman EA, Hays TS. The *Drosophila* tctex-1 light chain is dispensable for essential cytoplasmic dynein functions but is required during spermatid differentiation. Mol. Biol. Cell. 2004;15:3005–14.

53. Hecht NB. Molecular mechanisms of male germ cell differentiation. BioEssays. 1998;20:555–61.

54. McDonough CE, Whittington E, Pitnick S, Dorus S. Proteomics of reproductive systems: Towards a molecular understanding of postmating, prezygotic reproductive barriers. J. Proteomics. 2016;135:26–37.

55. Baker MA, Hetherington L, Weinberg A, Naumovski N, Velkov T, Pelzing M, et al. Analysis of phosphopeptide changes as spermatozoa acquire functional competence in the epididymis demonstrates changes in the post-translational modification of Izumo1. J. Proteome Res. 2012;11:5252–64.

56. Platt MD, Salicioni AM, Hunt DF, Visconti PE. Use of differential isotopic labeling and mass spectrometry to analyze capacitation-associated changes in the phosphorylation status of mouse sperm proteins. J. Proteome Res. 2009;8:1431–40.

57. Kalebic N, Sorrentino S, Perlas E, Bolasco G, Martinez C, Heppenstall PA. αTAT1 is the major α-tubulin acetyltransferase in mice. Nat. Commun. 2013;4:1962

58. Singh ND, Koerich LB, Carvalho AB, Clark AG. Positive and Purifying Selection on the *Drosophila* Y Chromosome. Mol. Biol. Evol. 2014;31:2612–23.

59. Riley NM, Coon JJ. Phosphoproteomics in the age of rapid and deep proteome profiling. Anal. Chem. 2016;88:74–94.

60. Walters JR, Harrison RG. Combined EST and proteomic analysis identifies rapidly evolving seminal fluid proteins in *Heliconius* nutterflies. Mol. Biol. Evol. 2010;27:2000–13.

61. Walters JR, Harrison RG. Decoupling of rapid and adaptive evolution among seminal fluid proteins in *Heliconius* butterflies with divergent mating systems. Evolution. 2011;65:2855–71.

62. Swanson WJ, Vacquier VD. The rapid evolution of reproductive proteins. Nat. Rev. Genet. 2002;3:137–44.

63. Hahn MW, Han MV, Han S-G. Gene family evolution across 12 *Drosophila* genomes. PLoS Genet. 2007;3:e197.

64. Zhao L, Saelao P, Jones CD, Begun DJ. Origin and spread of *de novo* genes in *Drosophila melanogaster* Populations. Science. 2014;343:769–72.

65. Haerty W, Jagadeeshan S, Kulathinal RJ, Wong A, Ravi Ram K, Sirot LK, et al. Evolution in the fast lane: rapidly evolving sex-related Genes in Drosophila. Genetics. 2007;177:1321–35.

66. Vicens A, Lüke L, Roldan ERS. Proteins involved in motility and sperm-egg interaction evolve more rapidly in mouse spermatozoa. PloS One. 2014;9:e91302.

67. Dorus S, Wasbrough ER, Busby J, Wilkin EC, Karr TL. Sperm proteomics reveals intensified selection on mouse sperm membrane and acrosome genes. Mol. Biol. Evol. 2010;27:1235–46.

68. Dean MD, Good JM, Nachman MW. Adaptive evolution of proteins secreted during sperm maturation: An analysis of the mouse epididymal transcriptome. Mol. Biol. Evol. 2008;25:383–92.

